# Electron cryo-tomography of vestibular hair-cell stereocilia

**DOI:** 10.1101/496513

**Authors:** Zoltan Metlagel, Jocelyn F. Krey, Junha Song, Mark F. Swift, William J. Tivol, Rachel A. Dumont, Jasmine Thai, Alex Chang, Helia Seifikar, Niels Volkmann, Dorit Hanein, Peter G. Barr-Gillespie, Manfred Auer

## Abstract

High-resolution imaging of hair-cell stereocilia of the inner ear has contributed substantially to our understanding of auditory and vestibular function. To provide three-dimensional views of the structure of stereocilia cytoskeleton and membranes, we developed a method for rapidly freezing unfixed stereocilia on electron microscopy grids, which allowed subsequent 3D imaging by electron cryo-tomography. Structures of stereocilia tips, shafts, and tapers were revealed, demonstrating that the actin paracrystal was not perfectly ordered. This sample-preparation and imaging procedure will allow for examination of structural features of stereocilia in a near-native state.

## Introduction

Hair cells of the inner ear detect minute forces arising from sound and head movements (Fettiplace and Kim, 2014). The mechanically sensitive structure of a hair cell is its hair bundle, an apical protrusion of ~100 actin-filled stereocilia (Roberts et al., 1988; Gillespie and Müller, 2009). Each stereocilium contains hundreds of actin filaments along most of its length, but that number dwindles near the base, where only a few dozen enter the cell. Stereocilia are arranged in a bundle in a highly patterned way, with a gradient of stereocilia height dictating the axis of mechanical sensitivity of the bundle. Stereocilia are connected together by a variety of links, with the tip link being of greatest interest. Tip links connect a stereocilium at its tip to the side of its tallest neighbor, and are aligned along the axis of sensitivity; tension in tip links opens transduction channels, the final step in the mechanical-to-electrical transduction that underlies auditory and vestibular function (Fettiplace and Kim, 2014).

Because of their highly regular structure, stereocilia are unusually amenable to investigation with ultrastructural imaging (Jacobs and Hudspeth, 1990; Hackney et al., 1993). Stereocilia structure usually is examined by transmission electron microscopy (TEM [^§^ Nonstandard abbreviations: cryo-ET, electron cryo-tomography; SEM, scanning electron microscopy; TEM, transmission electron microscopy; WT, wild type.]) of resin-embedded samples, with most samples having been chemically fixed, stained with heavy metals, dehydrated with alcohols, and embedded in resin prior to ultrathin sectioning. Each of those processing steps introduces the possibility of artifacts. An alternative to this approach is rapid-freeze, deep-etch imaging, where fixed or unfixed material is plunge-frozen and fractured, followed by etching to remove ice and observation in TEM (Hirokawa and Tilney, 1982; Hirokawa, 1986; Kachar et al., 2000). Coating with metal is required to obtain sufficient contrast, however, so freeze-etching still does not reveal native structure of the stereocilia cytoskeleton. Ultrarapid cryo-fixation of sensory epithelium improves preservation, but cannot fully eliminate possible sample artifacts—e.g., molecular aggregation and extraction—arising from heavy metal staining, dehydration and resin-infiltration protocols. Furthermore, stereocilia are ideally examined in a longitudinal or cross-sectional orientation, but samples are usually obtained in a slanted orientation, complicating ultrathin sectioning.

Electron cryo-tomography (cryo-ET) has emerged as a powerful method for examining macromolecular structures in cells (Baker et al., 2017; Oikonomou and Jensen, 2017; Hutchings and Zanetti, 2018). Samples frozen by rapid-freezing or high-pressure-freezing techniques retain their native structure, and tomography allows reconstruction of 3D structures within a slab of tissue. Cryo-ET is particularly useful for examination of native cytoskeletal structure (Jasnin et al., 2013; Turgay et al., 2017; McIntosh et al., 2018; Sun et al., 2018). We adapted stereocilia blotting methods (Neugebauer and Thurm, 1984; Shepherd et al., 1989; Hasson et al., 1997; Avenarius et al., 2017) to isolate a thin layer of stereocilia on electron microscopy grids. Stereocilia on grids were then plunge-frozen and imaged with cryo-ET. The resulting tomograms revealed key features of native stereocilia, including substantial imperfections in the paracrystalline arrangement of actin filaments in the stereocilia cytoskeletal core.

## Materials and Methods

### Preparation of electron microscopy grids

Tissue was blotted on lacey carbon film on 200-300 mesh gold or copper grids (Electron Microscopy Sciences). Grids were glow-discharged in a PELCO easyGlow 91000 (Ted Pella), and 15 nm gold fiducials (Sigma-Aldrich) were added at 1:4-1:10 dilution. Prior to dissection, grids were incubated with a 5 μl drop of 1 mg/ml poly-L-lysine on the sample side at room temperature, and the grids were covered to minimize evaporation; after at least 20 min, the poly-L-lysine solution was blotted off, and grids were allowed to dry for 2 hr. Coated grids could be stored for 2 days before use. Grids were glow-discharged again 30 min prior to blotting.

### Isolation of stereocilia on grids

Stereocilia isolation was similar to that previously reported (Neugebauer and Thurm, 1984). For wild-type C57BL/6 (WT) or plastin 1 knockout (*Pls1^-/-^*) mice, utricles were dissected from P21-P28 animals in Leibovitz L-15 media supplemented with 5 mM HEPES; otoconia were removed using an eyelash. Bullfrog sacculi were prepared for stereocilia blotting essentially as described (Shepherd et al., 1990). For mice or bullfrogs, organs were washed by transferring to a new dish of dissection buffer. To avoid disturbing the poly-L-lysine coating, the grid with its coated side up was placed into the dish by angling the edge of grid into the meniscus. The organ was then placed with the epithelium side down onto the center of the grid; gentle pressure was applied to the back of the organ with forceps in order to flatten it against the grid. The organ was then peeled away, leaving stereocilia attached to the grid.

### Plunge-freezing

After removing the tissue from the grid, the grid was carefully removed from the solution using an anti-capillary forceps and transferred to the manual plunger or Vitrobot. The remaining solution on the grid was then quickly blotted away by placing a small piece of filter paper at the edge of the grid interface with the forceps, and allowing most of the remaining solution on the grid and in between the forceps blades to wick away. A small aliquot of saline (4 μl) was added back, then manual blotting was carried out with Whatman #1 filter paper for 5-10 sec. The second blotting step allowed for better control of ice thickness, as the amount of solution left on the grids after dissection was highly variable and distinguishable by eye. Grids were plunge-frozen immediately after stereocilia isolation, and as quickly as possible after inner-ear dissection. Although most plunging was done on a home-made manual plunger (Lawrence Berkeley National Lab) (Comolli et al., 2012), a Vitrobot (ThermoFisher) was also used with similar success. When plunging was done in the Vitrobot, humidity was set to 70%, and the chamber was allowed to equilibrate 20-30 sec prior to loading the grid. The chamber door was left open and the humidifier was turned off during the majority of the dissection to prevent the filter paper becoming soggy and losing its ability to wick away solution. Grids were then stored under liquid nitrogen until imaging in the electron microscope.

### Electron cryo-tomography

Single-axis tilt series were generally collected to the full range allowed by the goniometer before the grid bar came into view, typically from +/−60° to 65° in 1.5-2° increments. Low-dose conditions were used (total dose of 80-100 electrons/Å^2^) on a Krios TEM (ThermoFisher) operated at 300 kV with a nominal defocus of −3.5 to −4.5 μm. Images were recorded on a Falcon 2 camera in integration mode at a pixel size of 0.47 to 0.59 nm. Reconstruction of all tomograms was done with IMOD (Kremer et al., 1996), using the gold fiducials for tilt series alignment and either weighted back-projection reconstructions or the SIRT reconstruction method provided by Tomo3D (Agulleiro and Fernandez, 2011). No obvious differences were observed between the two reconstruction methods. Cryo-tomograms were binned by 2 for reconstruction. Fiducials that were visible through the entire tilt series (typically 5-15) were selected by hand, and tracking of individual markers was checked and adjusted manually as necessary. Fiducials that could not be tracked accurately throughout the tilt series were ignored. The aligned tilt series was reconstructed with Tomo3d using the simultaneous iterative reconstruction technique (SIRT) (Agulleiro and Fernandez, 2011). The resulting tomograms were filtered with Priism (Chen et al., 1992), using five iterations (median) of the bilateral or median filters with a kernel size of 3 to improve contrast.

## Results

### Cryo-preservation of individual whole-mount stereocilia

Most previous two- and three-dimensional studies of hair-cell stereocilia using TEM have relied on ultrathin sections of resin-embedded tissue samples. Despite improved preservation due to high-pressure freezing and freeze-substitution, 3D tomographic imaging of the actin paracrystal in stereocilia (Shin et al., 2013) indicated that actin filaments and actin-actin crosslinkers were irregularly arranged throughout the imaged region. The ultrastructure of resin-embedded samples may have been compromised, however, by the addition of cryo-protectants prior to high-pressure freezing or by the lengthy freeze-substitution, resin-infiltration, embedding, and polymerization steps. We therefore sought to develop an alternative sample-preparation approach, preferably resulting in stereocilia samples that could be studied in an unstained, frozen-hydrated (vitrified) state by cryo-electron tomography, and that were therefore as close as possible to their native structure.

Building on a stereocilia blotting technique reported previously (Neugebauer and Thurm, 1984), we captured intact stereocilia onto an electron microscopy grid by bringing a freshly dissected sensory epithelium in the dissection buffer in contact with the grid and its lacey-carbon support film, which had been coated with poly-L-lysine. The epithelium was removed, leaving stereocilia adhering to the grid. Grids were removed from the solution, dissection buffer was wicked away, and samples were either chemically fixed (Fig. 1A) or plunge-frozen in liquid ethane (Figs. 2–4).

**Figure 1.**
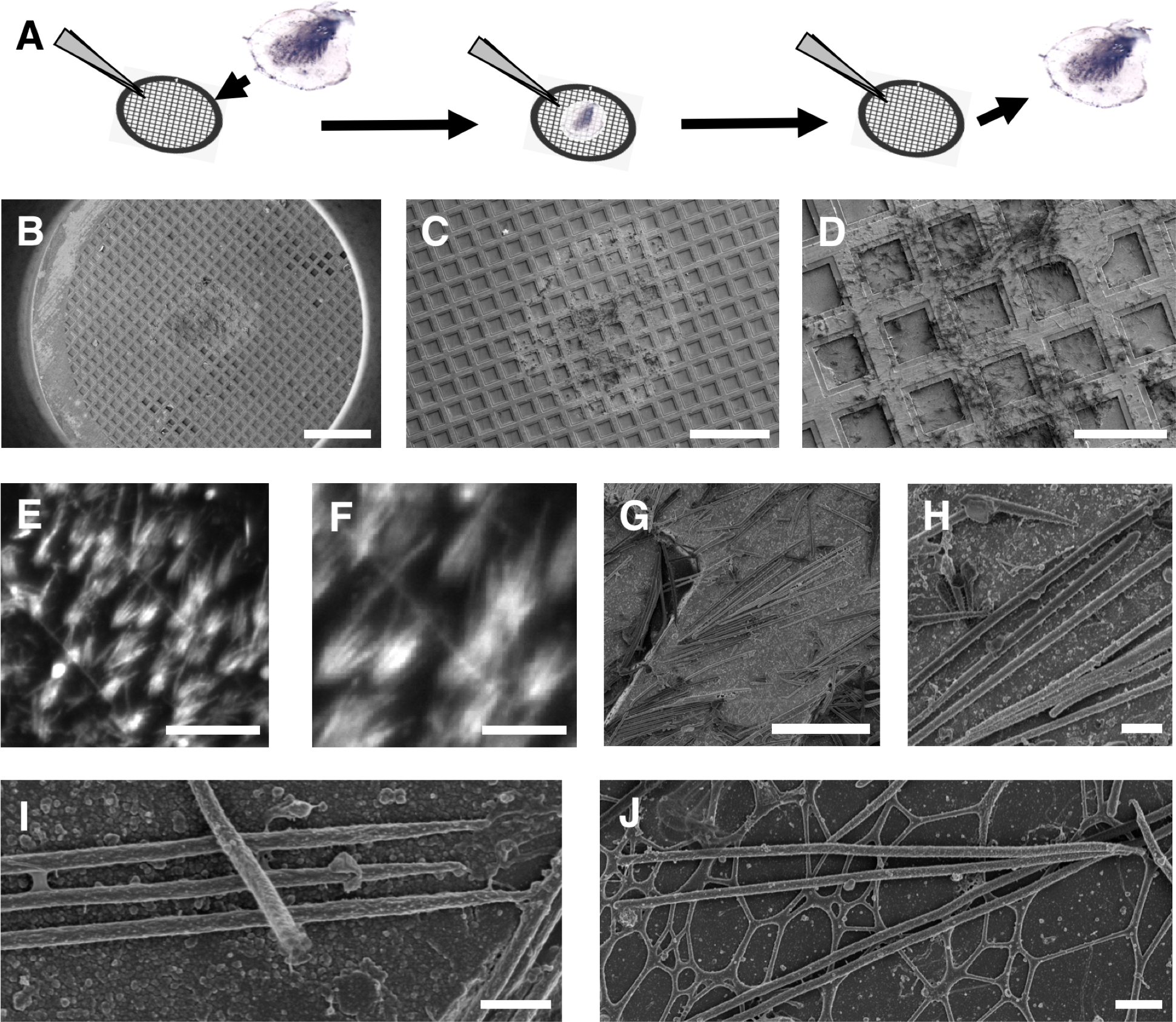
Stereocilia blotting from tissue surface onto electron microscopy support film. (**A**) Schematic view of stereocilia blotting. Sensory epithelia are brought briefly into contact with the electron microscopy support film in dissection buffer solution, resulting in the transfer of hair bundles and stereocilia onto the EM grid support film. (**B**) Overview of the entire electron microscope grid imaged by SEM. Note the imprint of the tissue at the center. Scale bar = 500 μm. (**C-D**) Close-up views of the center region of the grid imaged by SEM. Scale bars = 300 μm (C) and 100 μm (D). (**E-F**) Hair bundles labeled with Alexa 568-phalloidin. Scale bars = 50 μm (E) and 20 μm (F). (**G-H)** Low and intermediate resolution SEM images of stereocilia blotted into the grid bars. Scale bars = 10 μm (G) and 1 μm (H). (I) Higher resolution imaging of three adjacent stereocilia blotted onto an EM grid. Note that another stereocilium lays on top of the three-stereocilia monolayer. Scale bar = 1 μm. (**J**) Four stereocilia deposited onto the support film spanning the grid squares. Note that some stereocilia appear straight, whereas others show significant deviations from the original straight rod shape. Scale bar = 1 μm.

**Figure 2.**
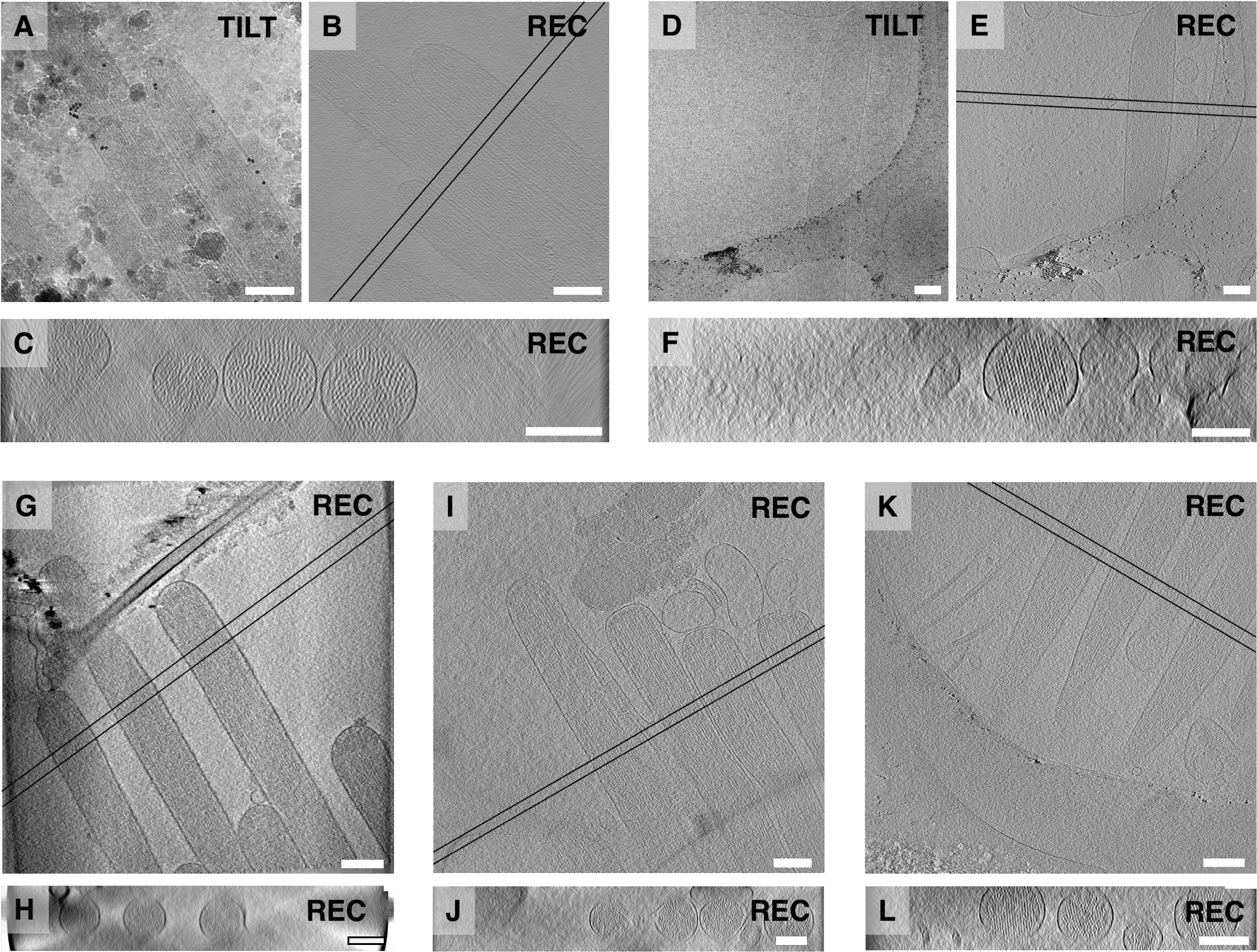
Tilt series and 3D reconstruction of stereocilia. **(A-C**) WT mouse utricle stereocilia shaft and tip region. (**D-F**) *Pls1*^-/-^ mouse utricle stereocilia shaft and tip region. (**G,H**) WT mouse utricle stereocilia shaft and tip region. (**I,J**) Frog sacculus shaft and tip region. (**K,L**) *Pls1*^-/-^ mouse utricle taper and rootlet region. (A) and (D) are zero-degree tilt series projection views; (B), (E), (G), (I), and (K) are central, longitudinal ~1 nm slices through the 3D reconstructions; (C), (F), (H), (J), and (L) are ~50 nm averaged cross-sectional views of stereocilia between the two parallel lines shown in (B), (E), (G), (I), and (K). Note the effect of the missing wedge on the 3D reconstruction as evidenced by anisotropic resolution in the z-direction (cross sections) compared to the x-y directions (longitudinal views). All scale bars = 200 nm.

We first characterized the isolation procedure by using scanning electron microscopy (SEM) to examine chemically fixed stereocilia on blotted grids. Stereocilia are shown in Fig. 1B-D at increasing magnification, with the ~1 mm imprint of the epithelium being readily visible in low-power views (Fig. 1B-C). Stereocilia were found scattered across the grid’s support film (Fig. 1D-J). Epifluorescence light microscopy of Alexa 568-phalloidin labeled stereocilia revealed a splayed, hair-bundle-like organization of stereocilia (Fig. 1E-F). High-magnification SEM imaging showed the stereocilia in more detail (Fig. 1G-J). In many cases, parallel stereocilia in close proximity formed a single layer (Fig. 1H-J), although two or more layers were often found (Fig. 1I). Most stereocilia appeared straight along their entire length and showed readily distinguishable tip and taper regions on either end of the shaft region. Other stereocilia showed significant deviations from linearity, which suggested physical damage to the stereocilia. Only well-preserved, straight stereocilia were used for actin-core analysis.

### Electron cryo-tomography of stereocilia

Using vitrified samples, we collected single-axis tomographic tilt series of young adult mouse utricle (Fig. 2A-H,K-L) and adult bullfrog saccule stereocilia (Fig. 2I-J). Mouse utricle samples included those dissected from plastin 1 knockout mutants (*Pls1*^-/-^), which in contrast to WT utricles, have hexagonally-packed actin filaments in their stereocilia (Krey et al., 2016). Low-dose projection images at zero-degree tilt are shown for WT (Fig. 2A) and *Pls1*^-/-^ (Fig. 2D) mouse stereocilia. Central sections through the corresponding 3D reconstructions are shown in longitudinal (Fig. 2B,E) and cross-sectional (Fig. 2C,F) orientations at positions indicated by two black parallel lines. Longitudinal views are single slices of ~1 nm thickness, whereas each cross-section view represents an average projection of a ~50 nm slab. Additional reconstructions of tip and shaft regions of WT mouse (longitudinal view in Fig. 2G) and bullfrog (Fig. 2I) stereocilia, as well as taper and rootlet regions of *Pls1*^-/-^ mouse stereocilia (Fig. 2K), are shown in the corresponding cross-sectional views (Fig. 2H,J,L). Shaft regions of mouse stereocilia were typically ~250 nm; bullfrog stereocilia shafts were 175-300 nm. Larger stereocilia diameter, thick vitrified ice and crystalline ice contaminations (Fig. 2A) are not ideal for tomographic in-depth analysis and lead to low feature contrast in 3D reconstructed volumes, as can be seen when comparing Fig. 2B with Fig. 2G. We focused our study on samples from the mouse utricle. In order to assess actin-filament organization, we calculated 50 nm cross-sectional averages (Fig. 2C,F,H,J,L). When viewed in cross section, most stereocilia appeared round, although stereocilia occasionally appeared flattened with the number of actin filaments parallel to the grid surface being significantly higher than the number of actin filaments perpendicular.

Actin paracrystals appeared better ordered in the *Pls1*^-/-^ data compared to those in WT data, in agreement with results from resin-embedded samples (Krey et al., 2016). We measured stereocilia diameters from 3D reconstructions, which allow us to confirm that stereocilia were not flattened. We had a larger dataset of *Pls1*^-/-^ stereocilia, which averaged 286 ± 49 nm (mean ± SD; n=21). Eight WT stereocilia had a diameter of 228 ± 45 nm (p=0.008). While this result was unexpected, as extensive characterization of *Pls1*^-/-^ stereocilia showed that they had a reduced diameter as compared to WT (Krey et al., 2016), the sample size was very small. Due to the missing wedge artifact, actin and membrane signals were strongest in the center and faded towards the top and the bottom of the tomographic reconstructions. We found a continuous membrane seal around the end of the taper region (Fig. 2K) at the site where stereocilia had been anchored in the cuticular plate prior to blotting of the hair bundles, similar to that shown previously with a different stereocilia isolation procedure (Gillespie and Hudspeth, 1991).

### Cryo-tomograms of tip, shaft, and taper regions

As shown in Fig. 3, we calculated 10 nm thin slab averages, corresponding to a single layer of actin filaments, of tip and shaft regions (Fig. 3A-H) and taper/rootlet region (Fig. 3I,J). Tips of bullfrog stereocilia were either rounded (Fig. 3A) or flattened (Fig. 3B), consistent with previous studies with resin-embedded samples (Jacobs and Hudspeth, 1990). Fig. 3C-E constitute mouse WT and Fig. 3F-J constitute *Pls1*^-/-^ tip/shaft and taper region data sets, respectively. For mouse samples, we found the tips to be mostly round, however, on occasion, we found a cone-shaped structure (Fig. 3F). For most data sets, actin filaments could be traced to the immediate vicinity of the tip membrane with a small gap of ~10-15 nm, except for one data set where we found a ~100 nm thin layer of what appears to be filamentous material oriented perpendicular to the actin bundle axis (Fig. 3H). In the shaft region, the actin filaments appeared well organized, with a smooth membrane in close proximity to the actin core (Fig. 3G-H). Occasionally the plasma membrane appeared detached from the actin core (Fig. 3C).

**Figure 3.**
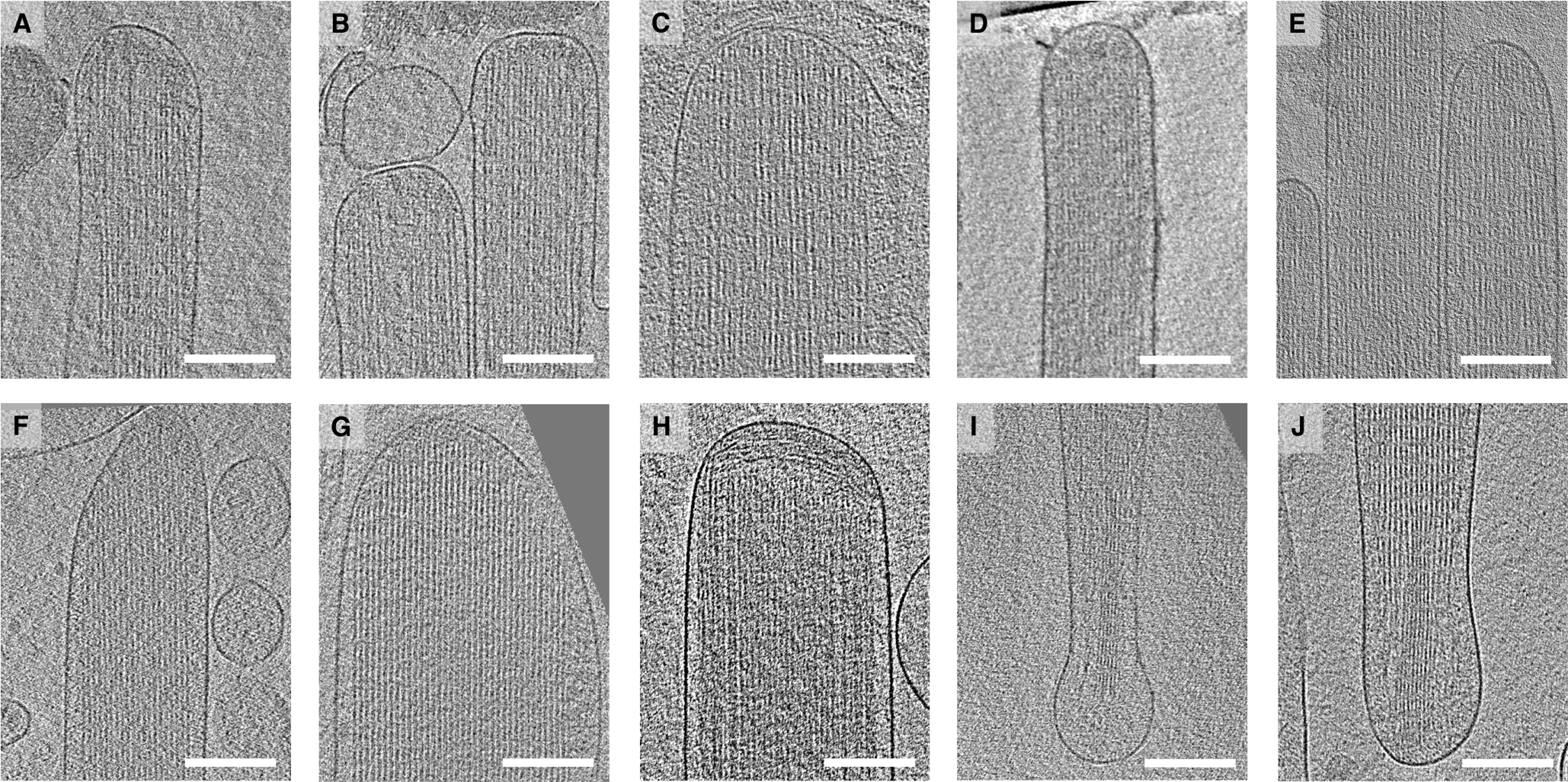
Tomograms of tip, shaft, and taper stereocilia regions. **(A-B**) Central slice through stereocilia tip region of frog sacculus. (**C-E**) Central slice through stereocilia tip region of wildtype mouse utricle. (**F-H**) Central slice through stereocilia shaft region of *Pls1*^-/-^ mouse utricle; note the presence of filamentous material near the tip in distinctly different orientation. (**I-J**) central slice through taper region of stereocilia from a *Pls1*^-/-^ mouse utricle. All scale bars = 200 nm.

In the taper region (Fig. 3I-J), actin filaments appeared to continue from the shaft region into the taper and rootlet regions. Interestingly, while the rootlet region in osmicated resin-embedded samples typically appeared considerably darker due to osmium binding (Furness et al., 2008), filaments did not appear notably different from that of the shaft region in our tomograms, although the filament packing appeared to be tighter (e.g., Fig. 3J). Individual filaments were readily distinguishable throughout the entire region.

The actin-actin spacing in these cryopreserved stereocilia was considerably larger than in conventionally processed samples (Krey et al., 2016). Examining one representative stereocilium each from WT and *Pls1*^-/-^ mice, we also found that that the actin-actin spacing was similar between genotypes, unlike what has been previously reported (Krey et al., 2016) (Fig. 4). WT actin-actin spacing was 12.5 ± 2.1 nm (mean ± SD; n=851), while *Pls1*^-/-^ spacing was 12.7 ± 1.3 nm (mean ± SD; n=925). The order in the *Pls1*^-/-^ stereocilia actin core was particularly apparent from this analysis (compare Fig. 4H to Fig. 4E).

**Figure 4.**
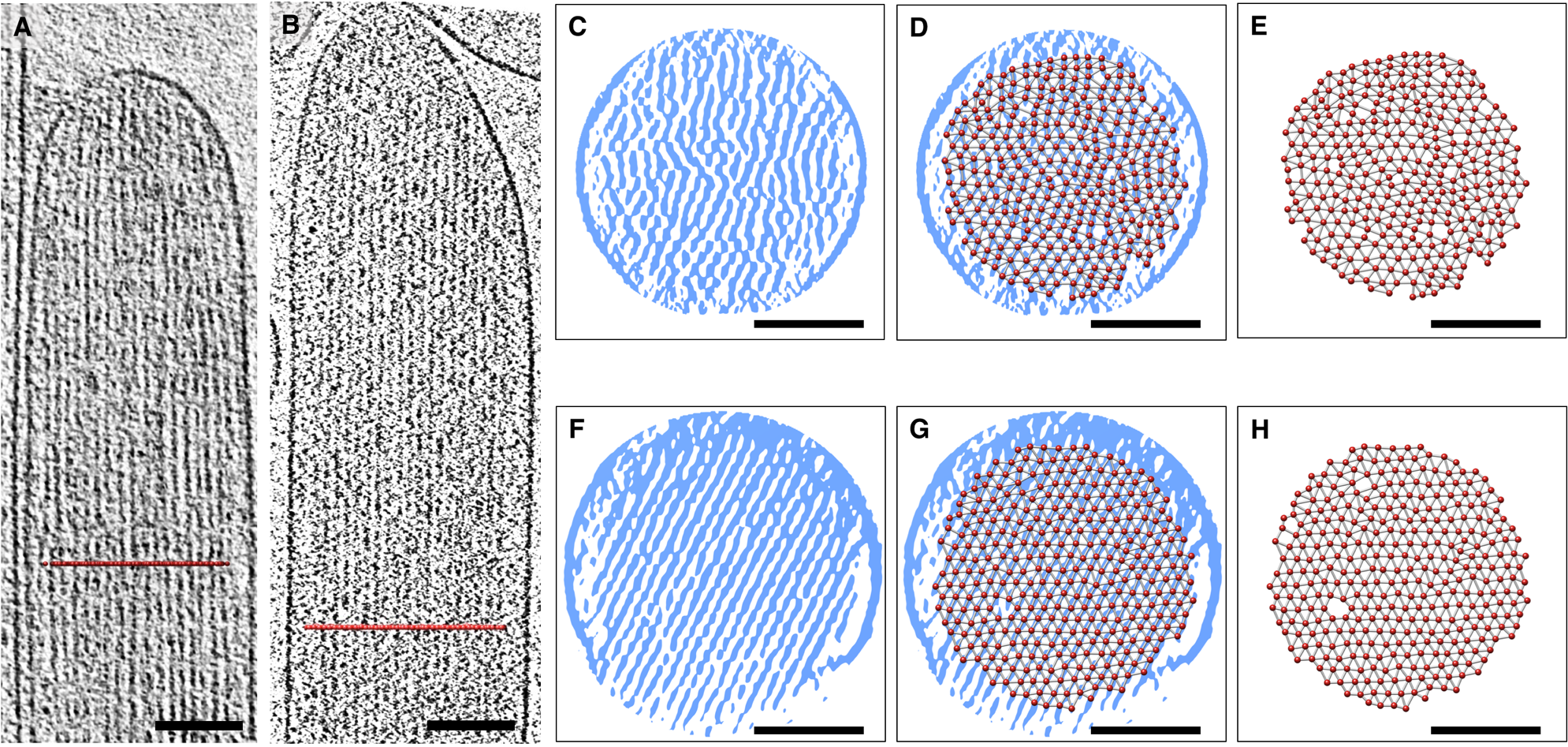
Analysis of actin-actin spacing. **(A-B**) Overview of a wildtype and *Pls1*^-/-^ stereocilia, respectively, shown as a ~10 nm slab of 3D tomogram. Colored lines indicate position of transverse section used for measuring actin-actin spacing. (**C,F**) Central slices through WT and *Pls1*^-/-^ stereocilia. (D,G) Central slices with actin filaments indicated with red circles. (**E,H**) Actin filaments are indicated by red circles and measurements between actin filaments (not position of crosslinkers) are indicated by lines. All scale bar = 100 nm.

We also observed a range of anomalies, including fusion of the membrane of adjacent stereocilia, stereocilia deformations, actin-core deformations, as well as membrane detachment in the tip and shaft regions. Whether such structures existed prior to blotting or were an artifact of stereocilia blotting onto the grid surface remains unclear.

### Actin filament order in stereocilia

To estimate the quality of our 3D reconstructions and their suitability for possible higher resolution reconstruction methods that exploit symmetry, such as the helical symmetry of the actin filaments or the expected hexagonal packing of the actin paracrystal for the *Pls1*^-/-^ (Krey et al., 2016), we performed fast Fourier transform (FFT) analysis of mouse WT (Fig. 5A) and *Pls1*^-/-^ (Fig. 5B) subregions of the tomographic reconstructions. While results are reported from 10 nm slabs, we found that analysis using smaller (6 or 8 nm) or larger (50 nm) slabs gave similar results. We used the Slicer tool in IMOD for creating a 10 nm slab containing approximately one actin filament layer, based on previous findings using high-pressure frozen and fixed samples (Krey et al., 2016). We oriented the slabs manually and calculated the corresponding Fourier transform from a projection of the slab approximately in the direction of the beam (Z-projection). Subvolumes of 512 x 512 pixels are shown as grey-scale density maps (Fig. 5C,F,I); we also used false-color representation, with blue for higher densities and yellow for lower densities (Fig. 5D,G,J), which emphasized actin filaments and the membrane. We examined three different regions of the stereocilia that were of particular interest: (1) the tip region (Fig 5C-E), which is expected to contain scaffolding for the transduction channel apparatus; (2) the shaft region ~1 μm below the stereocilia tip in WT (Fig. 5F-H), and (3) a shaft region just below the tip in *Pls1*^-/-^ (Fig. 5I-K). In some of our best samples, we see layer lines corresponding to well-established spacing of actin monomers within filaments, as well as row lines corresponding to actin-actin spacing, which is set by actin crosslinkers. These features suggest that our methods preserve well the long-range order of actin filaments, as well as protein structure, as is generally expected with vitrified samples. Improving further upon data collection and processing methods, such as recording data using electron-counting detectors with motion correction algorithms, could potentially allow us to gain a glimpse of the stereociliary protein machinery at near-atomic resolutions by applying subtomogram averaging or symmetry-based reconstruction methods. Despite all efforts to optimize orientation, however, imperfections of the actin paracrystal were evident, (Fig. 5C,D,F,G,I,J). FFT of the subregions are shown in Fig. 5E,H,K. Nonetheless, the presence of layer lines typical of actin filaments from in vitro experiments (Sukow and DeRosier, 1998; Volkmann et al., 2001; Sukow and DeRosier, 2003) in some of our best datasets suggests that our sample preparation procedure preserved the actin filament structure to near-atomic resolution.

**Figure 5.**
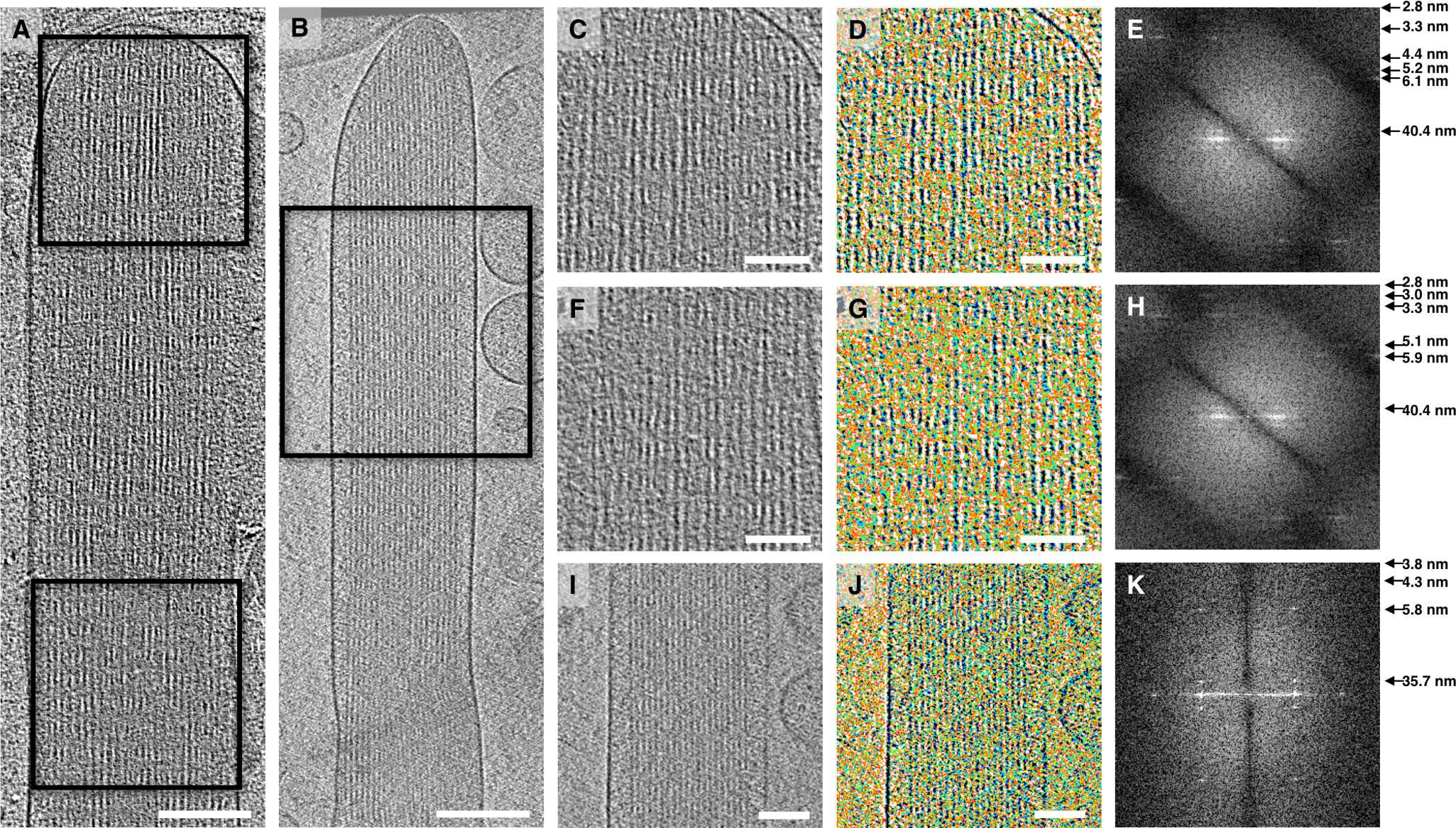
Fast Fourier transform analysis of single actin layer in a 3D tomogram. **(A-B**) Overview of a wildtype and *Pls1*^-/-^ stereocilia, respectively, shown as a ~50 nm slab of 3D tomogram. Scale bars = 200 nm. (**C**) Close-up view of the tip region shown in A. (**D**) False-color representation of (C), respectively, with the false-color assignment emphasizing the presence of ordered and disordered regions in stereocilia single actin-layers. (**E**) Fast Fourier Transform analysis of original map shown in (C) in IMOD, showing diffraction spots to 3.2 nm resolution. (**F**) Close-up view of the shaft region shown in (A). (**G**) False-color representation of F. (**H**) FFT analysis of (C), showing diffraction spots to 3.2 nm resolution. (**I**) Close-up view of the shaft region shown in (B). (**J**) False-color representation of **I. (K**) Fourier analysis of C, showing diffraction spots to 4.3 nm resolution. For (C), (D), (F), (G), (I), and (J), scale bars = 100 nm, density map is 10 nm slab average.

## Discussion

By blotting stereocilia onto the support film of the electron microscope grid, we have adapted a sample preparation procedure that allows plunge-freezing and cryo-tomographic imaging of frozen-hydrated stereocilia. Blotting was carried out using poly-L-lysine-coated surfaces, which have been used previously for imaging isolated stereocilia (Neugebauer and Thurm, 1984; Shepherd et al., 1989; Hasson et al., 1997; Avenarius et al., 2017). Blotted stereocilia on grids were readily visualized by fluorescence labeling, which emphasized that proteins retained their native states, or by SEM, which permitted visualization of many stereocilia in large fields of view.

Ultrarapid plunge-freezing of stereocilia immobilizes all macromolecules within milliseconds in a close-to-native state. Such vitrified, unstained stereocilia were of ideal size and shape for electron tomographic analysis, and different regions of the stereocilia could be targeted for data collection. Cross-sectional views of the electron tomographic 3D volumes revealed that most stereocilia retained their cylindrical shape over their entire lengths of 2-15 μm. In mouse preparations, actin filaments showed clear hexagonal packing for *Pls1*^-/-^ stereocilia, consistent with a previous report (Krey et al., 2016), but packing was less regular for WT stereocilia. The anisotropy of the data in the Z- versus the X,Y-direction, which is a direct result of limitations of data collection geometry, can be seen in stereocilia membrane cross-sectional views.

Stereocilia membranes typically reseal after isolation from epithelial tissue (Gillespie and Hudspeth, 1991), and here we show that the membranes often formed a bulb-like structure at the taper end. We suggest that the membrane bending energy (Evans, 1974) prevents a tight radius of curvature around the taper and rootlet, particularly when the membrane detaches from the cytoskeleton.

The spacing between actin filaments was considerably larger than anticipated from earlier experiments using conventionally fixed stereocilia (Krey et al., 2016). Actin-actin spacing was greater both in WT (9.7 nm in(Krey et al., 2016), 12.5 nm here) and *Pls1*^-/-^ WT (7.9 and 12.7 nm). The spacing reported here is similar to that found in other actin arrays cross-linked with PLS1 (Volkmann et al., 2001) and is consistent with the near-native preservation offered by cryopreservation.

Our method for stereocilia isolation and vitrification provides unprecedented preservation, which will allow future segmentation and model-building of the actin-filament core with its crosslinkers and actin-membrane connectors for the tip, shaft, taper and rootlet regions of stereocilia. Notably, Fourier analysis suggests a preservation of structural information of macromolecular complexes in stereocilia at 4-6 nm resolution. Data shown here were recorded in the integrated charge-detection mode, and thus were not subject to motion-correction; with new direct electron-counting detectors (Kuijper et al., 2015) and phase-plate technology (Danev and Baumeister, 2017), data quality is expected improve further. The methodology reported here demonstrates further progress towards developing a high-resolution model of a critical organelle.

## Acknowledgements

MA acknowledges support by National Institutes of Health (NIH) grant P01 GM051487. PGBG was supported by NIH grants R01 DC011034 and P30 DC005983; DH was supported by NIH grant R01 GM115972; and NV was supported by NIH grant P01 GM121203. NIH grants S10 OD012372 (DH) and P01 GM098412-S1 (DH) funded the purchase of the Titan Krios transmission electron microscope and Falcon II direct used for some data acquisition in this study. We received support from the OHSU Multiscale Microscopy Center, where additional data were acquired using a Titan Krios electron microscope.

